# FORGE: multivariate calculation of gene-wide p-values from Genome-Wide Association Studies

**DOI:** 10.1101/023648

**Authors:** Inti Pedroso, Michael R Barnes, Anbarasu Lourdusamy, Ammar Al-Chalabi, Gerome Breen

## Abstract

Genome-wide association studies (GWAS) have proven a valuable tool to explore the genetic basis of many traits. However, many GWAS lack statistical power and the commonly used single-point analysis method needs to be complemented to enhance power and interpretation. Multivariate region or gene-wide association are an alternative, allowing for identification of disease genes in a manner more robust to allelic heterogeneity. Gene-based association also facilitates systems biology analyses by generating a single p-value per gene. We have designed and implemented FORGE, a software suite which implements a range of methods for the combination of p-values for the individual genetic variants within a gene or genomic region. The software can be used with summary statistics (marker ids and p-values) and accepts as input the result file formats of commonly used genetic association software. When applied to a study of Crohn’s disease susceptibility, it identified all genes found by single SNP analysis and additional genes identified by large independent metaanalysis. FORGE p-values on gene-set analyses highlighted association with the Jak-STAT and cytokine signalling pathways, both previously associated with CD. We highlight the software’s main features, its future development directions and provide a comparison with alternative available software tools. FORGE can be freely accessed at https://github.com/inti/FORGE.

## Introduction

Genome-wide association studies (GWAS) have led to the discovery of hundreds of replicated variants associated with diverse human phenotypes ^1^. However, most GWAS have low statistical power to detect true effects due to the low effect-sizes of common risk alleles and the need for robust genome-wide significance thresholds, to allow for multiple testing, e.g. 7.2 x 10^−8^ ^2^. Multivariate analytical strategies, such as gene-wide association, are an attractive alternative to single SNP approaches. They have the capability to allow for allelic heterogeneity (independent associated alleles in the same region) (e.g. ^3^), result in fewer tests genome-wide ^4^ and provide gene p-values that can be used with gene-set analysis methods ^5^. Increasing evidence suggest that allelic heterogeneity is a common feature of the genetic architecture of complex traits and that gene-set methods can improve the interpretation and statistical power of GWAS by using prior biological knowledge ^6-8^.

Numerous gene-wide association methods have been proposed, for example ^9-13,13,14^. The simplest form consists of correcting the minimum p-value in the region or gene by its number of single nucleotide polymorphisms (SNPs), e.g., a Bonferroni correction or a measure of effective number of tests ^15,16^. This approach ignores possible allelic heterogeneity, i.e., independent potentially associated alleles. Multivariate methods are an alternative but they can require computationally expensive simulations to derive significance if the test statistic’s null distribution is unknown, as may be due to the correlation between genetic markers.

Currently available software to perform gene-based association include: i) VEGAS that implements a simulation-based strategy to estimate significance ^13^; ii) PLINK with an implementation of Hoteling’s T2-statistics and Makambi’s modified Fisher’s test statistic ^17^, --T2 and --set-screen options respectively; and iii) GATES that implements a modified Simes test ^14^. PLINK also allows users to perform SNP-set analyses, e.g., all SNPs within genes of a biological pathway, allowing for gene-set analyses. Analysis of GWAS using gene-set analysis has been reviewed recently elsewhere, e.g., ^5,6^, and we provide a representative list of available tools and studies in Supplementary Table 1 and 2, respectively.

Here we describe FORGE, a software suit to perform gene-wide and gene-set analyses of GWAS. It provides routines for the calculation of four gene-wide association methods and two gene-set analysis strategies. FORGE represents an extension and complements parallel independent efforts, such as VEGAS ^13^ or KGG ^14^,^18^. In addition, several utility programs are distributed with FORGE allowing users to: i) map SNP to Genes using the Ensembl human genome annotation; ii) parse different gene-set files; and iii) calculate meta-analysis statistics for gene and genesets analyses results when studies carried out on multiple data-sets.

## Methods

### Gene-wide statistics

#### 1. Sidak’s correction on minimum p-value

Consider a set of *m* SNPs (**M**) and their association p-values (**P**) with a trait, for which the aim is to calculate a combined association test statistic for **M**. The simplest strategy is to consider the minimum p-value among **P** as **M**’s evidence for association using Sidak’s correction ^19^ with an estimate of the number of effective tests within the gene, *p*_*sidak*_ = 1 −(1 − *p*_*raw*_)^*k*^, where *p*_*raw*_ is the minimum p-value in **P** and *k* is the effective number of SNPs tested calculated with the method of ^16^.

#### 2. Modified Fisher’s method to combine correlated p-values

In order to obtain a combined test (**T**) for **M** taking the correlation among the genetic markers into account, we use the method originally derived by Brown (1975) with the modifications proposed by Kost and McDermott ^20^ and Makambi ^17^, leading to the chi-square test *T* = 0.5 × 0 ×*M _F,m_*, with *ν* degrees of freedom, where 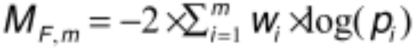 is the weighted version of the Fisher’s method, *p*_i_ is the p-value of *i*th marker and *w*_*i*_ are weights greater than zero that sum to one. The degrees of freedom are *ν* = 8/var(*M*_*F,m*_) with 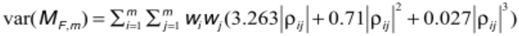, where *ρ*_*ij*_ is the correlation between the P_i_ and P_j_.

#### 3. Fixed-effect z-score statistic

We can also calculate 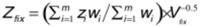(Huedo-Medina *et al.*, 2006), where *z*_*i*_ are the p-values transformed to z-scores using the standard normal distribution inverse cumulative distribution function (c.d.f.) and V_*fix*_ is the variance of the test. Using the approximation of the multivariate-normal distribution 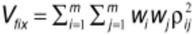.

#### 1. Random-effect z-score statistic

A random-effects estimate is given by 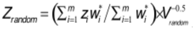, with variance 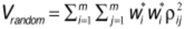 and weights equal to 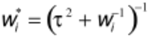 which are adjusted with the heterogeneity measure 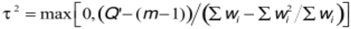. In calculating *τ*^2^ one would normally use Cochran’s heterogeneity statistics 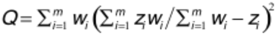, which is an approximately distributed chi-square variable with *m*-1 degrees of freedom ^21^. To account for the correlation among the genetic markers *τ*^2^ is calculated using Q’, which is Q re-scaled into a chi-square variable with *m*-1 degrees of freedom. This is achieved by calculating Q’s tail probability using the modified Fisher’s method described above and then Q’ is the probability’s chi-square value from a chi-square distribution with *m*-1 degrees of freedom.

### Gene-set analysis

#### 1. SNP to gene-sets strategy

In this case we treat gene-sets as a large gene, i.e. map to it all SNPs of its genes and applied the statistics described above.

#### Gene-sets analysis with gene p-values

We implemented the methods described by Luo et al. ^22^ as following. GSA is performed by transforming the gene p-values into z-scores (using the standard distribution inverse c.d.f.) and combining the z-scores with 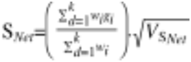 where *g*_*i*_ is the z-score of the *i*^th^ gene in the gene-set, *k* the number of genes in the gene-set and *V*_*SNet*_ is the variance-covariance matrix of the gene’s statistic, 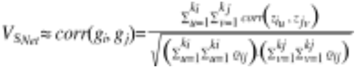. S_*Net*_ is formally a variable from a standard normal distribution and it significance can be estimated with normal distribution probability density function.

### Calculation of gene p-values

The gene-wide statistics describe above lead to asymptotic estimates of significance. We implemented the method described by Li et al. ^14^ to approximate the correlation between the p-values by the correlation between the SNPs, i.e. allelic LD or Pearson’s correlation. In addition to these asymptotic strategy we implemented routines to calculate gene p-values using the simulation-based strategy of Liu et al. ^13^. Although this strategy is slower than asymptotic methods, its p-values are well correlated with empirical estimates ^13^. We refer the reader to Liu et al. ^13^ for details of the strategy and to Supplementary Material for description of our implementation.

### Analysis of the WTCCC Crohn’s disease GWAS

We obtained summary statistics of the Crohn’s disease GWAS from the EGA website with formal data access permission of the WTCCC Data Access Committee. Quality control (QC) performed QC by excluding samples as indicated in the files provided by the WTCCC and SNPs with missingness ≥ 1%, minor allele frequency (MAF) ≤ 1% Hardy-Weinberg equilibrium (HWE) p-value ≤ 1 x 10^−3^ and p-value ≤ 1 x 10^−5^ in controls and cases, respectively, as previously described ^23^. We also obtained genotype data of the bipolar disorder GWAS and performed QC as follow: i) poor quality samples and SNPs were removed as indicated in the files distributed by the WTCCC; and ii) SNPs were removed using the same criteria used for the Crohn’s disease summary statistics. SNP association were performed with a logistic regression as implemented on PLINK ^24^. Using both sets of summary statistics gene-wide statistics were calculated for approximately 19,550 protein-coding, long intergenic non-coding RNA and micro-RNA genes annotated in Ensembl version 59 and whose SNPs passed QC in the WTCCC studies. We mapped SNPs to genes if the SNP was within 20 kb of the annotated coordinates aiming to include 95% of potential eQTL loci ^25^. We approximated the correlation between test statistics as the correlation between the SNPs as measured by the Person’s correlation between the allele counts. We used a False Discovery Rate (FDR) < 0.1 ^26^ to perform multiple testing correction on the gene p-values.

We performed gene-set association with the WTCCC Crohn’s disease gene-wide p-values results using 5,384 gene-sets derived from the Human Protein Reference Database protein-protein interaction network (PPIN)^27^. The PPIN gene-sets were constructed with the following algorithm: subnetwork searches started from each node (seed node) in the PPIN and a subnetwork was defined by adding sequentially the direct neighbours of the subnetwork’s nodes (initially only the seed node). We allowed searches to go to a max of 5 interactions from the seed node and generate subnetworks of 2 to 200 nodes in size; each subnetwork was used as a gene-set. An FDR of 0.1 was used for multiple testing correction ^28^. Significant PPIN’s genes were analysed by assessing their over-representation among biological categories reported on KEGG ^29^ and GO databases ^30^. Significance of the overlap was calculated with binomial statistics.

## Results and Discussion

Figure 1 presents the GWAS analyses implemented in FORGE. GWAS summary statistics are used to calculate gene-set p-values either by mapping SNP p-values to gene-sets directly or by calculating gene-based p-values as an intermediate step. Calculation of gene p-values provides an additional layer of analysis that can facilitate integration of GWAS with results from other “omics” technologies that also generate a single statistic per gene, e.g., gene-expression studies. This integration can also be performed at the level of gene-sets.

**Figure 1.**
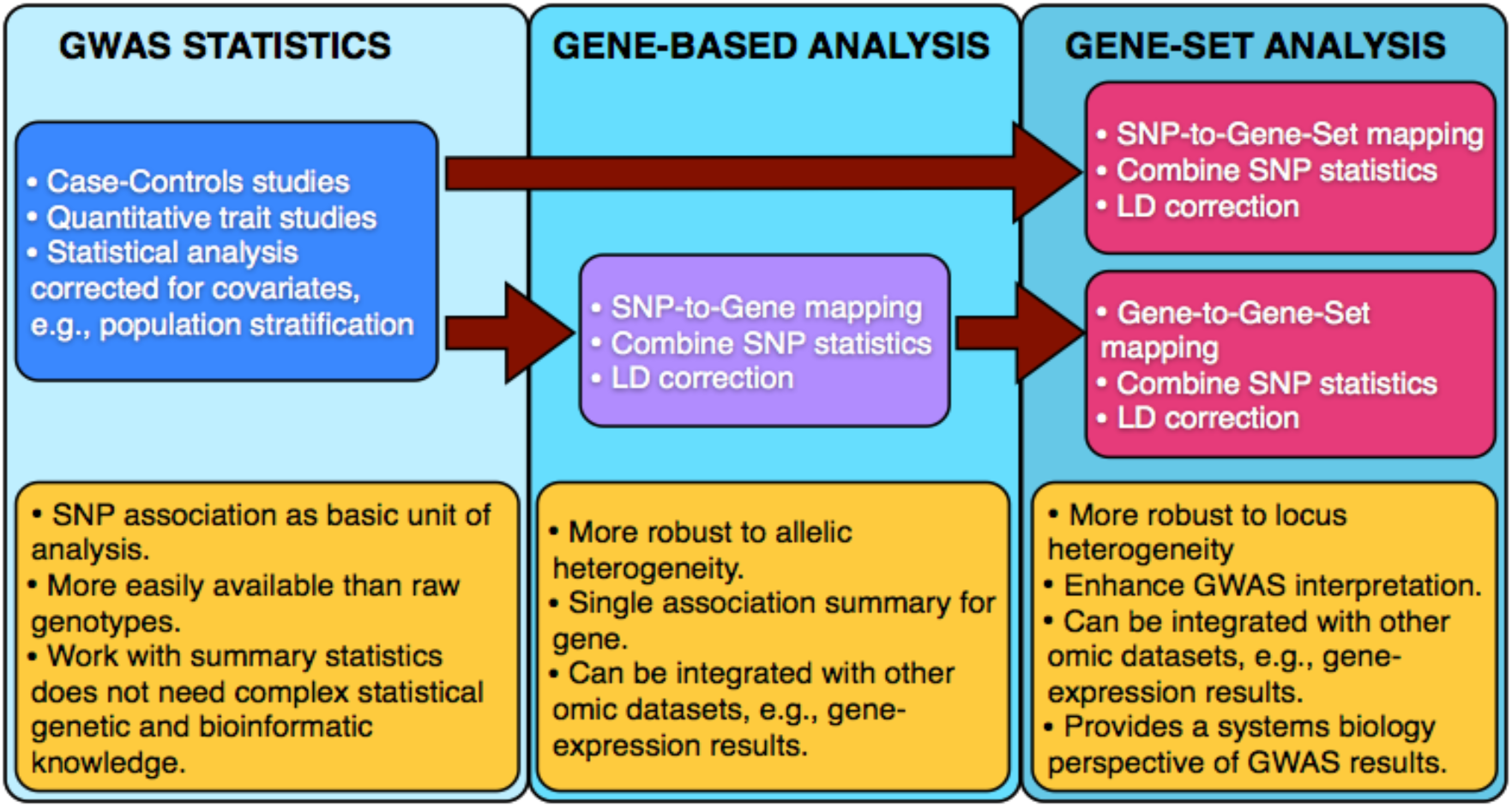
GWAS analyses implemented on FORGE. A FORGE analysis starts with GWAS summary statistics. It is possible to calculate gene-set statistics directly from SNP association p-values or by calculating gene p-values as an intermediate step. In all steps FORGE introduces corrections for the LD to avoid inflation of the statistics. Yellow boxes present the advantages and potential of each GWAS result level.

Liu et al. ^13^ introduced a simulation-based gene-wide association strategy that provides gene p-values with very good agreement with those obtained by phenotype permutations. For all gene-wide association methods we implemented this strategy as a way to estimate significance. This approach requires computation of correlations between SNP genotypes. It is desirable to use genotypes from a reference population, e.g. HapMap project samples, because it enables the use of summary statistics in the absence of each study’s genotype data. Using the WTCCC bipolar disorder GWAS we calculated gene p-values using the WTCCC genotypes and the HapMap Phase 3 CEU samples. The computing time decreased approximately linearly with the sample size, i.e., the computing time is reduced by ∼50 times when using the 165 HapMap Phase 3 CEU samples compared with using the full WTCCC Crohn’s disease dataset (5000 samples) (not shown). Also there was high correlation (>0.99) between the gene p-values calculated with both sets of genotypes (not shown), as may have been expected due to the Caucasian origin of both samples. There was high correlation between all gene-wide p-values (minimum correlation > 0.7) and the correlations between multivariate methods were higher (minimum correlation > 0.95) (Figure 2). Importantly, the distribution of gene p-values for the bipolar disorder GWAS did not present over-dispersion (Figure 3), in agreement with the idea that this particular GWAS was underpowered to detect the true genetic effects ^31^. Finally, we compared the results of the fixed and random-effects models and found small differences but only on relatively small genes, i.e., approximately < 45 SNPs (Figure 4). For larger genes the statistical heterogeneity measure I^2^ is rarely different from zero. This pattern is to be expected since only a small fraction of genetic variation is thought to be associated with a phenotype, so in larger genes most of the evidence will point to lack of association and statistics will be more homogeneous (low I^2^). Thus, our exploration of using a statistical heterogeneity measure to tackle genetic heterogeneity suggest it is hardly effective in moderate to large genes. Application of other statistical techniques may provide better results, for example ^32,33^.

**Figure 2.**
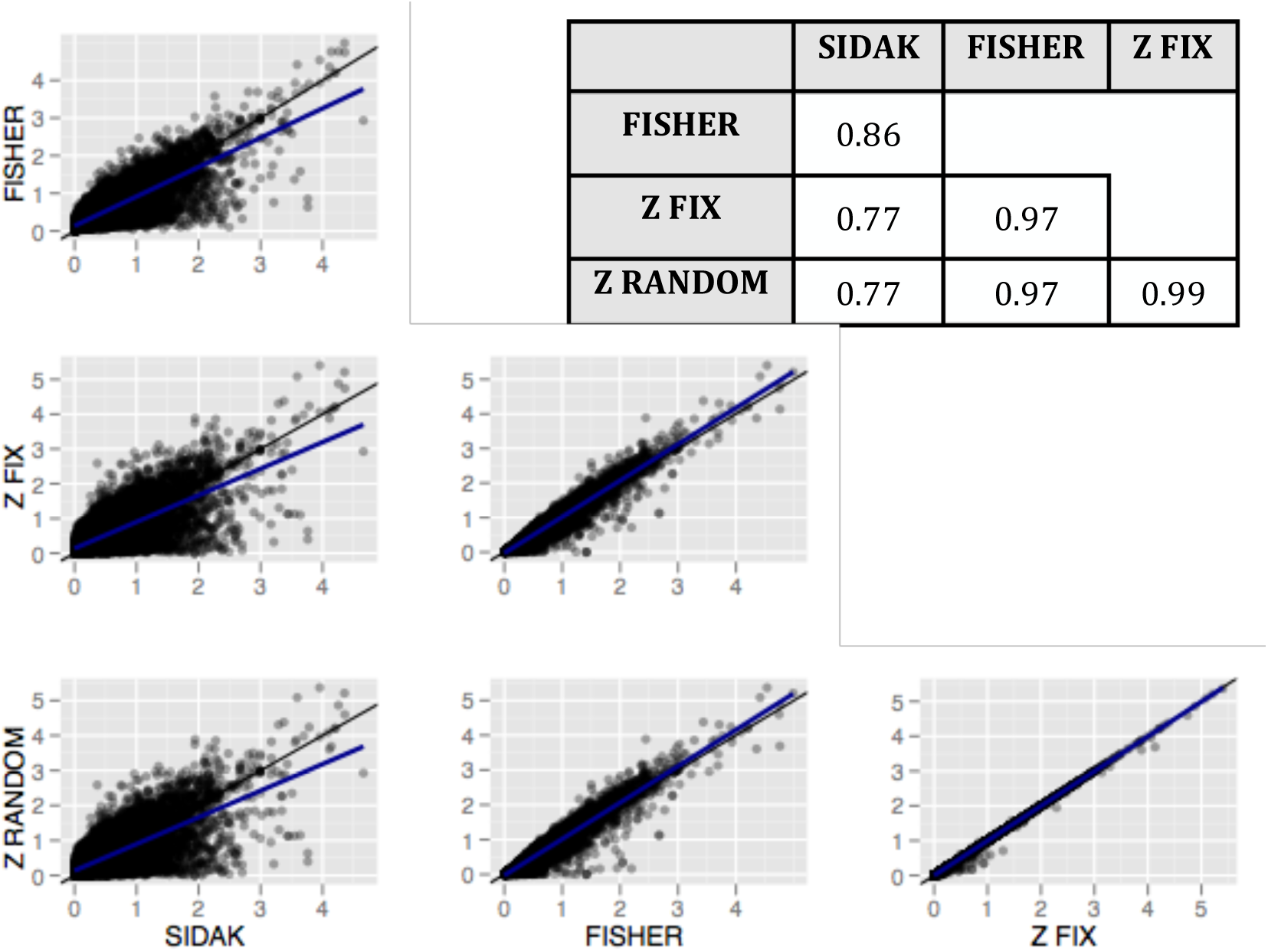
Correlation between gene-based p-values. Black diagonal line represents the 1-to-1 correlation. Blue line is a trend calculated with a linear model. Correlation estimates between the methods are indicated on the table.

**Figure 3.**
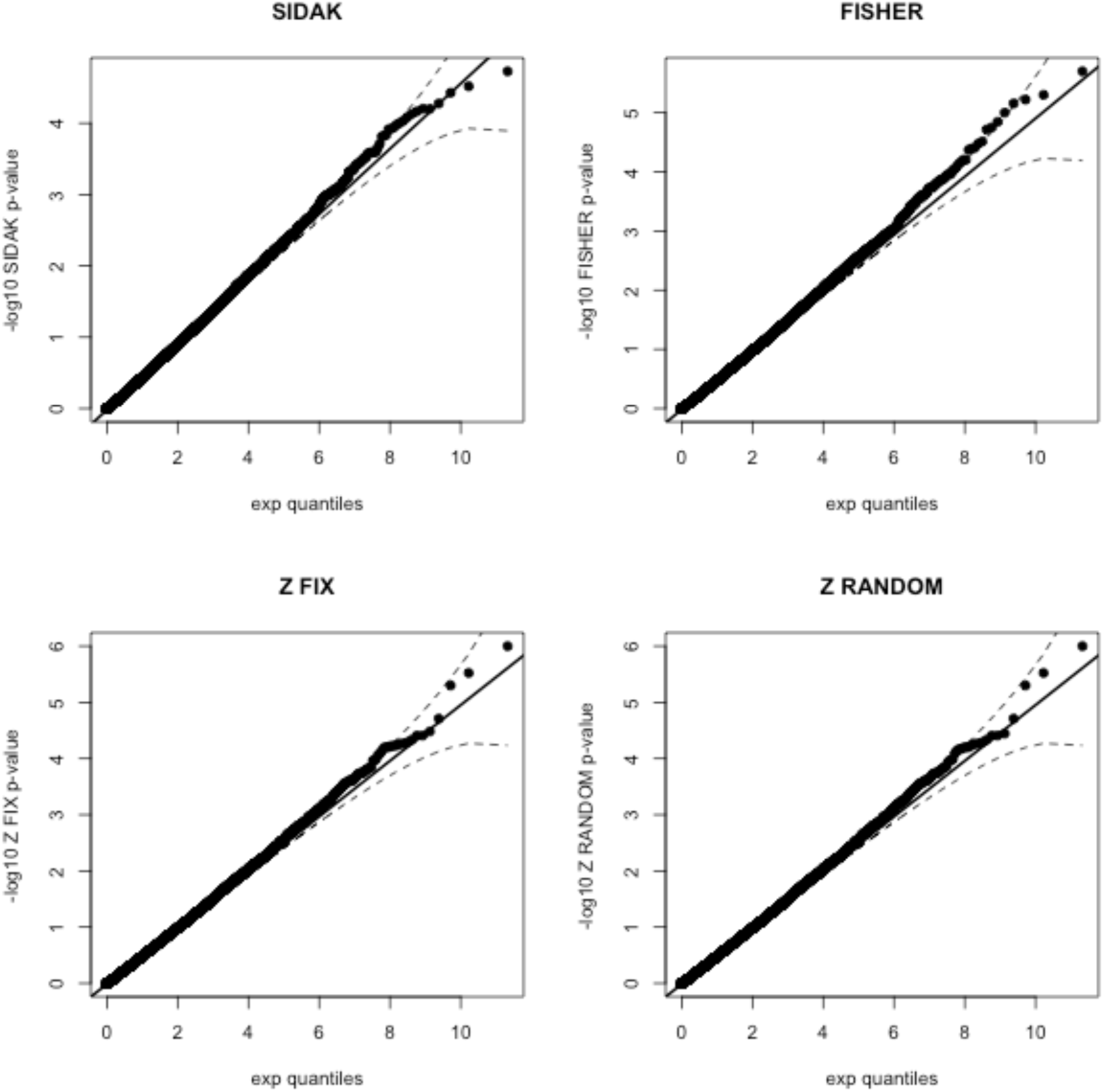
Quantile-quantile plots of simulation-based gene p-values. Plotted is the expected (x-axis) against the observed (y-axis) – log10 of the gene p-values. Dotted line marks the 95 % confidence interval.

**Figure 4.**
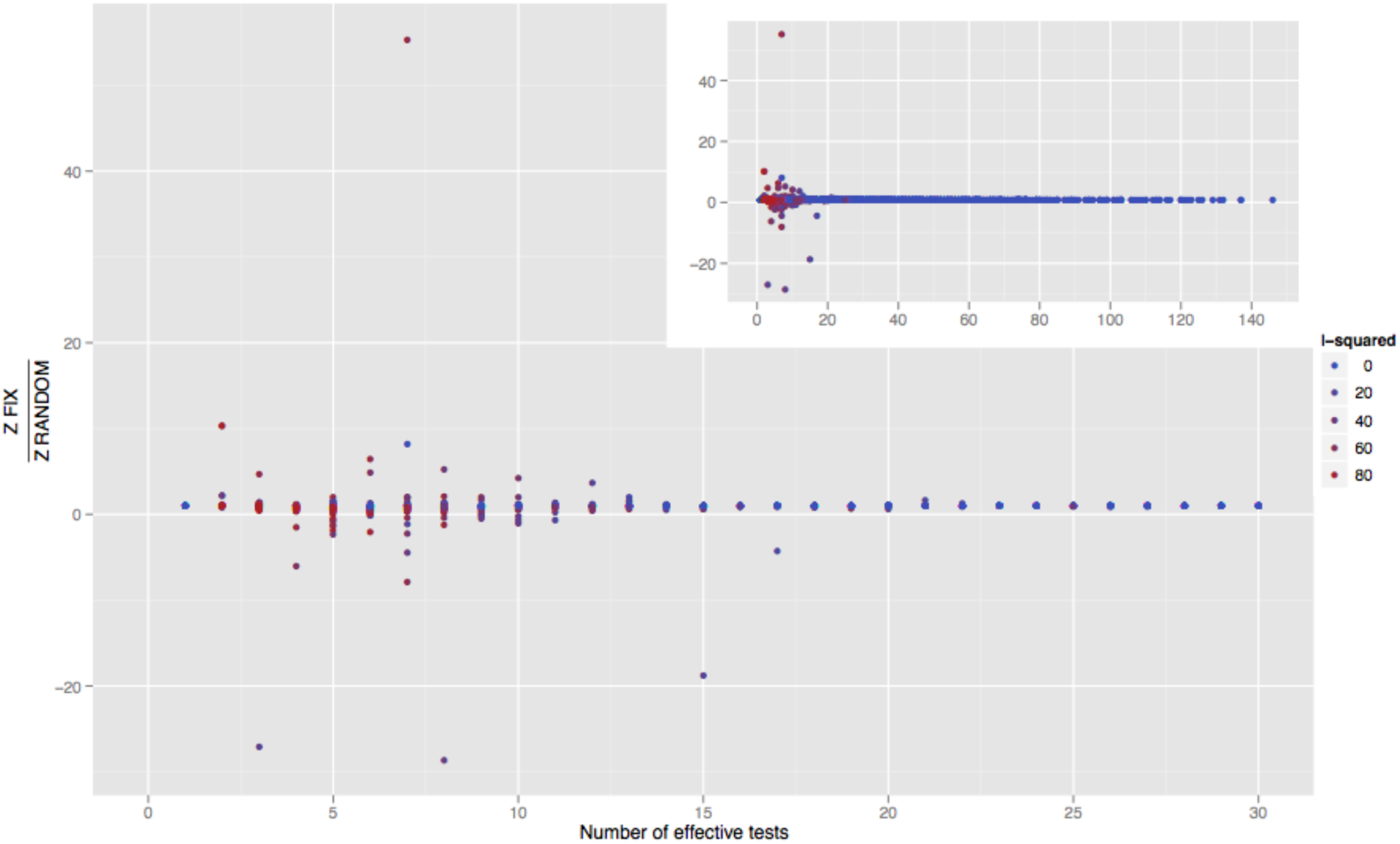
Comparison of fix and random-effects estimates. Plotted for each genes (N ∼20000) is the log10 of the ratio of fix and random-effects gene p-values. Insert (top right corner) presents data for the complete range of number of effective tests. Points are coloured (see figure legend) as the gene statistical heterogeneity measure I2, which has range from 0 (no evidence of heterogeneity) to 100% (maximum evidence of heterogeneity).

### Comparison with other software

Table 1 presents a comparison of FORGE with other software available to perform gene and gene-set analyses on GWAS. Compared with most other software, FORGE allows to perform both gene and gene-set analyses functionalities within the same software. In addition, it implements more analysis methods including asymptotic and simulation-based calculations. FORGE also reads all three major genotype formats used in GWAS and provides utilities to build SNP-to-gene mapping with updated versions of the human genome annotation. Finally, it Perl implementation makes it platform independent.

**Table 1.**
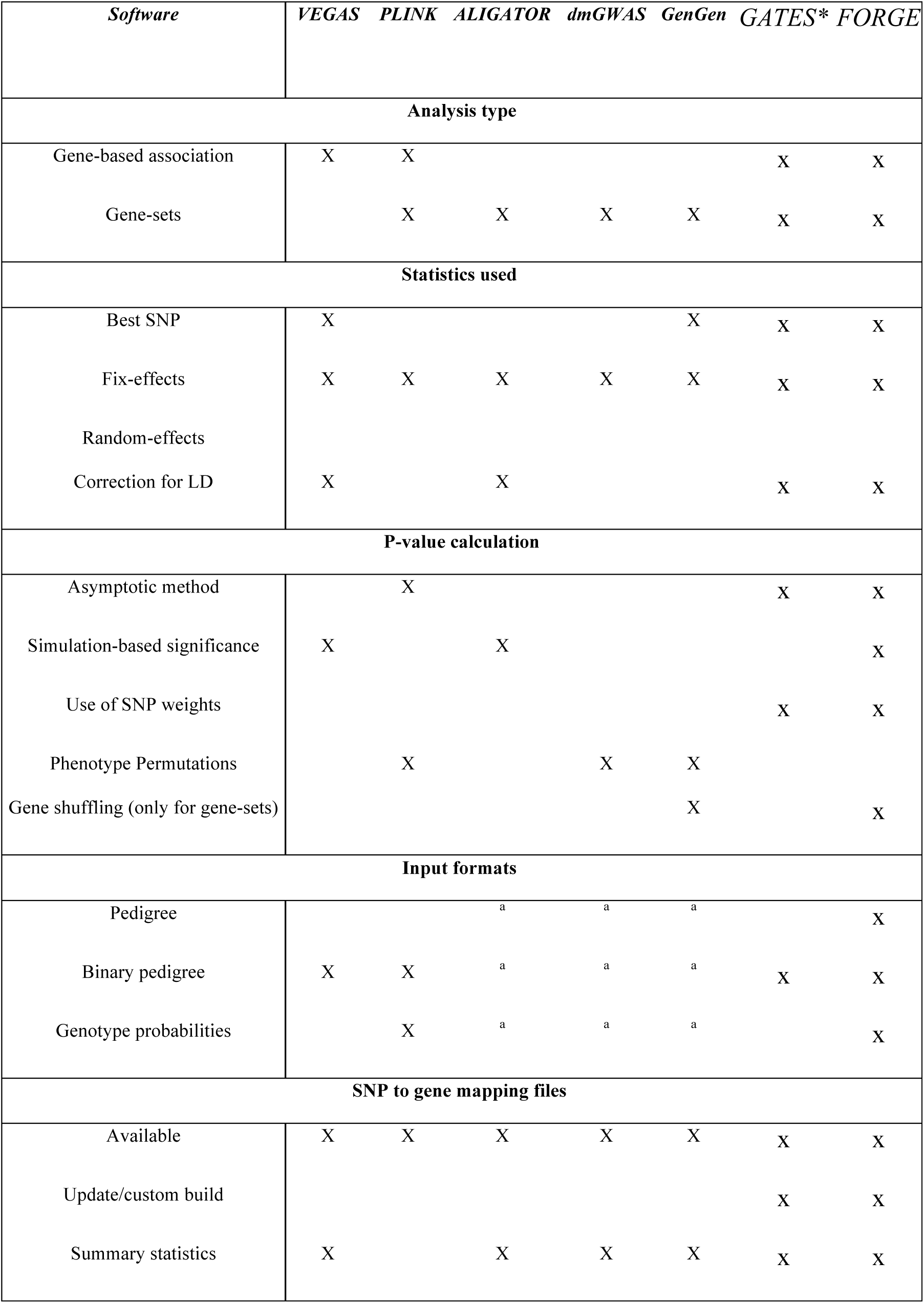
Comparison of FORGE with other software to perform gene and gene-set analyses of GWAS. ^a^ = method used by software does not need genotype files. * reported features correspond to the KGG software (http://bioinfo1.hku.hk:13080/kggweb/) where GATES has been made available.

### Case study: the WTCCC Crohn’s disease GWAS

Wang et al. ^6^ recently reviewed gene-set analyses results of GWAS of CD and we use these as a reference for compare results. FORGE gene-based association was performed using summary statistics and genotypes of the HapMap phase 3 CEU samples genotypes. Table 2 lists genes with FDR < 0.1 on the Z FIX gene-based association method. Genes mapped to significant SNPs (p-values < 5x10^−8^) were also identified by the gene-based association. Interestingly, in line with results of Liu et al.

**Table 2.** Genes with FDR < 0.1 from WTCCC CD GWAS. Z FIX p-value was calculated with the simulation-based strategy using up to 106 simulations. FDR = false discovery rate. a Genes within 20 kb of genome-wide significant SNPs from the CD GWAS meta-analysis reported by Franke et al. 37.

| Gene Symbol | Position | Uncorrected<br>Minimum p-value | Z FIX | FDR |
| --- | --- | --- | --- | --- |
| CYLD<sup>a</sup> | 17q12 | $4 \times 10^{-14}$ | $< 1 \times 10^{-6}$ | $1 \times 10^{-3}$ |
| ATG16L1<sup>a</sup> | 5q31.1 | $4 \times 10^{-13}$ | $< 1 \times 10^{-6}$ | $1 \times 10^{-3}$ |
| NKX2-3<sup>a</sup> | 16q12.1 | $1 \times 10^{-8}$ | $< 1 \times 10^{-6}$ | $1 \times 10^{-3}$ |
| IL23R<sup>a</sup> | 2q37.1 | $3 \times 10^{-12}$ | $< 1 \times 10^{-6}$ | $1 \times 10^{-3}$ |
| NOD2<sup>a</sup> | 3p21.31 | $3 \times 10^{-14}$ | $< 1 \times 10^{-6}$ | $1 \times 10^{-3}$ |
| C5orf56<sup>a</sup> | 2p13.1 | $1 \times 10^{-6}$ | $< 1 \times 10^{-6}$ | $1 \times 10^{-3}$ |
| IRGM<sup>a</sup> | 2p13.1 | $1 \times 10^{-7}$ | $< 1 \times 10^{-6}$ | $1 \times 10^{-3}$ |
| ZNF300<sup>a</sup> | 1q32.1 | $1 \times 10^{-7}$ | $2 \times 10^{-6}$ | 0.02 |
| PTPN2<sup>a</sup> | 10q24.2 | $1 \times 10^{-7}$ | $2 \times 10^{-6}$ | 0.02 |
| MST1<sup>a</sup> | 5q31.1 | $1 \times 10^{-6}$ | $3 \times 10^{-6}$ | 0.02 |
| ENSG00000249738 | 5q33.3 | $5 \times 10^{-5}$ | $1 \times 10^{-5}$ | 0.09 |
| ENSG00000221733<sup>a</sup> | 2p13.1 | $2 \times 10^{-11}$ | $2 \times 10^{-5}$ | 0.09 |
| APEH<sup>a</sup> | 5q33.1 | $1 \times 10^{-6}$ | $4 \times 10^{-6}$ | 0.09 |
| HLA-DQA2 | 2p13.1 | $5 \times 10^{-5}$ | $3 \times 10^{-5}$ | 0.09 |
| KIF21B<sup>a</sup> | 19q13.31 | $7 \times 10^{-5}$ | $3 \times 10^{-5}$ | 0.09 |
| CCL18 | 1p31.3 | $1 \times 10^{-4}$ | $5 \times 10^{-5}$ | 0.09 |
| TAP2 | 3p21.31 | $5 \times 10^{-5}$ | $5 \times 10^{-5}$ | 0.09 |
| HLA-DOB | 3p21.31 | $5 \times 10^{-5}$ | $5 \times 10^{-5}$ | 0.09 |
| ENSG00000250264 | 16q12.1 | $5 \times 10^{-5}$ | $5 \times 10^{-5}$ | 0.09 |
| RNF123<sup>a</sup> | 17q21.2 | $4 \times 10^{-5}$ | $5 \times 10^{-5}$ | 0.09 |
| ZNF283 | 8p23.1 | $1 \times 10^{-4}$ | $6 \times 10^{-5}$ | 0.09 |
| LOXL3 | 3p21.31 | $2 \times 10^{-4}$ | $6 \times 10^{-5}$ | 0.09 |
| DAG1 | 3p21.31 | $3 \times 10^{-5}$ | $7 \times 10^{-5}$ | 0.09 |
| IRF1 | 3p21.31 | $4 \times 10^{-6}$ | $7 \times 10^{-5}$ | 0.09 |
| P4HA2 | 18p11.21 | $4 \times 10^{-4}$ | $8 \times 10^{-5}$ | 0.09 |
| STAT3<sup>a</sup> | 5q31.1 | $2 \times 10^{-5}$ | $8 \times 10^{-5}$ | 0.09 |
| USP4 | 6p21.32 | $6 \times 10^{-5}$ | $8 \times 10^{-5}$ | 0.09 |
| ENSG00000245942 | 6p21.32 | $3 \times 10^{-4}$ | $8 \times 10^{-5}$ | 0.09 |
| GMPPB | 1p31.3 | $1 \times 10^{-4}$ | $9 \times 10^{-5}$ | 0.09 |
| C2orf65 | 6p21.32 | $2 \times 10^{-4}$ | $9 \times 10^{-5}$ | 0.09 |
| TNKS | 5q33.1 | $1 \times 10^{-4}$ | $9 \times 10^{-5}$ | 0.09 |
| HTRA2 | 6p21.32 | $2 \times 10^{-4}$ | $1 \times 10^{-4}$ | 0.09 |

^13^ and Li et al. ^14^, gene-based association results highlighted genes associated at a genome-wide significant level in other studies, in spite of having sub-threshold association on the GWAS analysed here. We used gene p-values calculated with the Z FIX method for gene set analyses (Table 3). Several significant associations were found among biological process with previous compelling evidence of involvement of Crohn’s disease pathology ^6^, for example: associations with gene-sets of the JakSTAT (hsa04630) and Cytokine-cytokine receptor interaction (hsa04060) signalling pathways.

**Table 3.**
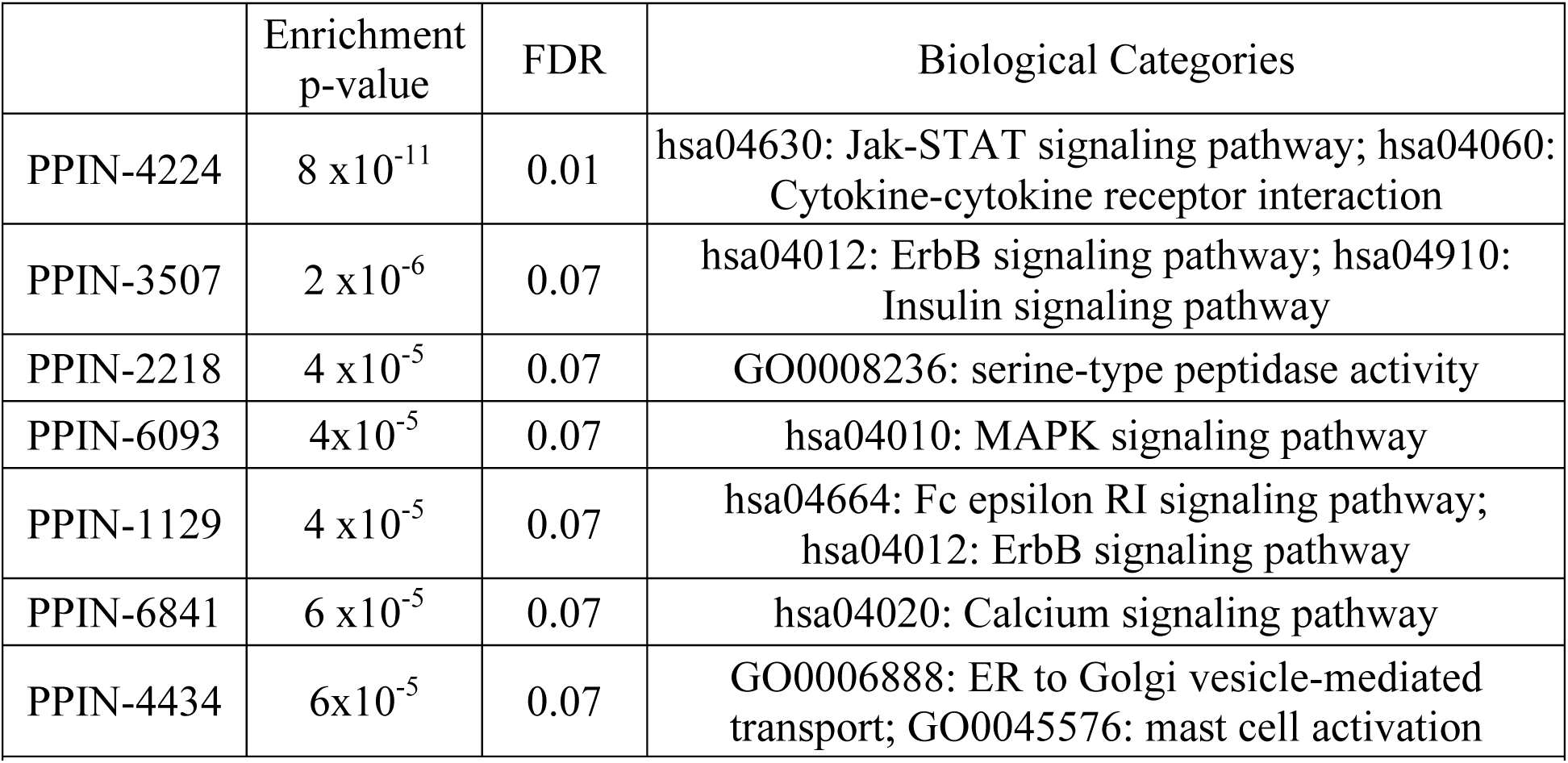
Protein-protein interaction networks with FDR < 0.1 on CD GWAS. Reported biological categories were over-represented among the subnetwork genes.

Overall, FORGE produces gene p-values that complement single SNP associations. Its gene p-values can be calculated from summary statistics and used with gene-set analysis methods to provide a systems biology perspective of a GWAS. Together, gene and gene-set association represent a complement to single SNP analyses by helping to interpret and extract information from GWAS.

## Availability and Future Directions

FORGE has been deposited in the public repository GitHub (https://github.com). Software code is updated using control version with GIT (http://git-scm.com) and users can access the latest stable or development versions at https://github.com/inti/FORGE. Current development is focused on a web-server version of FORGE to run on the 264 CPU computer cluster hosted at the NIHR BRC Centre for Mental Health (Institute of Psychiatry, KCL, UK). This interface will allow users to upload GWAS summary statistics and perform both gene and gene-set analysis online.

## Design and Implementation

We implemented these methods in a software suite called FORGE written in Perl (www.perl.com) using the PDL, PDL::Stats and PDL::LinearAlgebra libraries, all freely available at the Comprehensive Perl Archive Network (www.cpan.org). These libraries allow performing calculation on double mathematical precision with the General Scientific Library (GSL) and the LAPACK library. All major functions have been implemented as separate libraries, for example: i) GWAS_STATS.pm with statistical analysis routines; ii) GWAS_IO.pm to deal with file formats commonly used in GWAS studies; and iii) CovMatrix.pm implements method to calculate correlations and covariance matrices. This design allows new features to be developed in a modular fashion and the use of specific functions (e.g., reading binary genotype files) from software by the wider scientific community.

Three major programs were developed: i) forge.pl to perform gene and gene-set analyses; ii) gsa.pl implements additional routines to perform a gene-set analyses; and iii) meta_analysis.pl implements to combined results from independent studies.

### Main features

1. Input files: the program reads input and output files and genotype file formats of commonly used GWAS analysis software, e.g., genotype files in Pedigree, Pedigree Binary Format and SNP association files produced by PLINK ^24^. Geneset definitions are accepted in GMT format, described elsewhere (www.broadinstitute.org/cancer/software/gsea/wiki/index.php/Main_Page).
2. SNP-SNP correlations: three measures of SNP-SNP correlations are implemented: i) shrinkage correlation estimate described by Schafer and Strimmer ^34^; ii) LD; and iii) correlation among the test statistics as explained by Li et al. ^14^.
3. SNP-to-Gene annotations: pre-computed files with genetic variants mapped to genes up to 500 kb from gene coordinates are available at the FORGE website. A Perl script to update the annotation using the Ensembl API ^35^ is distributed as a utility of FORGE.
4. Additional features: i) user provided SNP weights, e.g. functional scores, can be used and will be re-scaled to sum to 1 within each gene; ii) genomic-control correction ^36^ can be automatically performed within the program; iii) analyses can be restricted to chromosomes, gene lists or gene types (e.g. protein coding or miRNA genes); iv) Affymetrix SNP identifiers are accepted and mapped to rsids internally; and vi) gene-sets for major databases like KEGG ^29^ or GO ^30^ are provided as well as those derived from the protein-protein interaction network (see Methods).
5. Documentation: Example files and tutorials are available on the software’s website.

## Acknowledgments

Funding: IP, GB and AAC thank funding from the NIHR Biomedical Research Centre for Mental Health at the South London and Maudsley NHS Foundation Trust and Institute of Psychiatry, Kings College London. GB and AAC thank the Medical Research Council. GB also thanks the Wellcome Trust, NARSAD and BBSRC for funding. AAC thanks The Motor Neurone Disease Association of Great Britain and Northern Ireland, ALS Therapy Alliance, Angel Fund and ALS Association for support.

